# Impact of methicillin resistance on virulence factor expression in *Staphylococcus aureus*: Insights from gene expression profiling

**DOI:** 10.1101/2024.10.19.619075

**Authors:** Charfeddine Gharsallah, Asma Ferjani, Mouna Ghariani, Sana Ferjani, Lamia Kanzari, Amel Rehaiem, Ahmed Fakhfakh, Ilhem Boutiba Ben Boubaker

## Abstract

*Staphylococcus aureus* is a major human pathogen causing various clinical infections and a leading cause of morbidity and mortality worldwide. S. aureus infections are problematic due to frequent antibiotic resistance, especially to methicillin. This study investigated 30 unduplicated S. aureus strains from clinical samples to establish a link between methicillin resistance and virulence factors.We detected and determined expression levels of the mecA gene, virulence genes (spdC, spA, atlA), and the RNAIII regulator using qRT-PCR. All virulence genes and the RNAIII regulator were detected in all strains. Phenotypic results showed only three strains (10%) were methicillin-resistant, while 12 (40%) carried the mecA gene. mecA-positive strains exhibited high expression of adhesion factors (spA) and biofilm formation factors (atlA), but low expression of the RNAIII regulator. The regulator’s expression was negatively correlated with mecA gene expression. Using a multilayer association network, we found a correlation between phenotypic methicillin resistance expression and mecA gene transcription in *S. aureus* mecA+. Understanding *S. aureus* virulence determinants will help develop anti-virulence strategies, especially given the lack of an anti-*S. aureus* vaccine and rising antibiotic resistance.

**Highlights:**

- **Complex interplay between methicillin resistance and virulence:** Our study unveils a complex interplay between methicillin resistance and the expression of virulence genes in Staphylococcus aureus clinical isolates.
- **Phenotypic and molecular correlation:** Phenotypic resistance to methicillin was observed in only 10% of the isolates, whereas 40% carried the *mecA* gene. Molecular analysis revealed distinct expression patterns, notably elevated *spA* and *atlA* expression, in *mecA*^+^ strains.
- **Negative correlation with *RNAIII*:** Our findings indicate a negative correlation between *RNAIII* regulator expression and the *mecA* gene in the same strains, shedding light on their regulatory relationship.
- **Multilayer association network:** Utilizing a multilayer association network, we established a correlation between phenotypic methicillin resistance and *mecA* gene transcription in *S. aureus mecA*^+^ strains.

## Introduction

*Staphylococcus aureus* causes a wide variety of diseases, ranging from moderately severe skin infections to life-threatening diseases [1]. It is a leading causative agent in pneumonia and other respiratory tract infections, surgical site, prosthetic joint and cardiovascular infections, as well as nosocomial bacteremia [2]. Recent developments have increased research efforts into unraveling *S. aureus* virulence mechanisms. These developments include, first and foremost, the rise in the early 2000s of community-associated (CA) methicillin-resistant *S. aureus* (MRSA), strains that combine methicillin resistance with high virulence potential in a previously unknown fashion [3]. This pathogen is characterized by its high capacity to express many virulence factors [4]. Surface proteins have many roles, especially adhesion, evasion of the immune response, and biofilm formation. In fact, *S. aureus* is an adept biofilm creator, which enhances its virulence capacity to establish persistent infections [5]. Studies have shown that *spdC*, a surface protein, was recently identified as a new virulence factor. It was shown that *spdC* inhibits the *WalK*R-dependent synthesis of peptidoglycan hydrolase [6]. The *atlA* and *SpdC* genes are involved in biofilm formation, and *spA*, encoding protein A, participates in adhesion and invasion in human tissue [6,7]. The expression of these three factors is controlled via genetic loci such as *agr, sar, sae* and *xpr*. The accessory gene regulator (*agr*) locus is the most important regulatory locus [8–10]. Infections with MRSA are accompanied by increased mortality, morbidity, and length of hospital stay compared to those caused by methicillin-sensitive *S. aureus* (MSSA) [11]. However, the presence of a broad spectrum of virulence genes in the genomes of MSSA strains could act as a potential source of infection. Thus, MSSA should be given the same attention as MRSA strains [12]. This study aimed to compare the genetic expression of *spA, spdC, atlA* and *RNAIII* genes in MRSA and MSSA clinical strains and to highlight the crucial role of *RNAIII* in the regulation of the expression of these virulence genes and the *mecA* gene.

## 1. Materials and Methods

### 1.1. Bacterial strains

Thirty nonreplicated clinical *S. aureus* strains isolated from different human specimens were collected between March 1 and August 31, 2020, in the Microbiology Laboratory of Charles Nicolle University Hospital of Tunis, a 1100-bed tertiary care facility.

### 1.2. Bacterial identification and antimicrobial susceptibility testing

*S. aureus* isolates were identified by conventional methods, including Gram staining, catalase, coagulase, deoxyribonuclease (DNase), and mannitol fermentation tests. When needed, biochemical identification with API-Staph (bioMérieux, Marcy l’Etoile, France) was used. Antimicrobial susceptibility testing was performed using the agar disk diffusion method, according to the European Committee of Antimicrobial Susceptibility Tests [13,14]. The following antimicrobial disks were tested: cefoxitin (30 μg), kanamycin (30 μg), gentamicin (10 μg), tobramycin (10 μg), tetracycline (30 μg), chloramphenicol (30 μg), ofloxacin (5 μg), trimethoprim-sulfamethoxazole (1.25/23.75 μg), erythromycin (15 μg), clindamycin (2 μg), pristinamycin (15 μg), linezolid (30 μg), teicoplanin (30 μg), vancomycin (10 μg), fosfomycin (5 μg), rifampin (5 μg) and fusidic acid (10 μg). For MRSA, the MICs of vancomycin and teicoplanin were determined by the broth microdilution method using the UMIC vancomycin/teicoplanin test (Biocentric, Bandol, France).

### 1.3. Genotypic characterization and quantitative real-time RT-PCR

#### 1.3.1. DNA extraction and PCR assays

The strains were grown on brain heart infusion (BHI) cultures at 37 °C overnight. Genomic DNA used as a target for PCR assays was extracted using a DNeasy Blood and Tissue kit (Qiagen, USA) according to the manufacturer’s instructions. Three virulence factor genes (*spA, atlA*, and *SpdC*), *RNAIII* of the *agr* system, and the *mecA* gene were detected by PCR as previously described [15–17]. The primers targeting the virulence genes (*spA, atlA, SpdC*), the ARNIIII regulator, and the *mecA* gene are shown in **Supporting Information Table S1**.

#### 1.3.2. RNA isolation and quantitative real-time PCR analysis

Total RNA was isolated using the RNeasy Mini kit (Qiagen, USA) according to the manufacturer’s instructions. RNA was quantified by an ND-1000 spectrophotometer (Nanodrop Technologies, USA). First-strand cDNA was synthesized from 250 ng of total RNA with reverse primers for the *SpA, AtlA, SpdC, RNAIII* and *mecA* genes and 200 U/L MMLV reverse transcriptase (Invitrogen, USA) according to the manufacturer’s instructions. The ABI 7500 sequence detection system (Applied Biosystems, USA) was used for quantitative real-time PCR (qPCR) under the following cycle conditions: 10 min at 95 °C followed by 40 cycles of 15 s at 95 °C and 1 min at 55-60 °C. The 16S rRNA gene was used as an internal reference gene [6]. PCRs were carried out in 96-well optical reaction plates (Applied Biosystems, USA). The reaction included 50 ng of cDNA sample as a template, 400 nM forward and reverse primers, and Igreen qPCR master Mix-Rox (BIOMATIK, USA). Relative quantification was performed by applying the 2 ^−ΔΔCt^ method [18].

#### 1.3.3. Biological network

Protein-protein interactions were retrieved from the STRING database (version 5.0) (http://string-db.org/). These interactions were filtered based on confidence scores, with a minimum interaction score of 0.400, thus generating a network of interactions by Cytoscape (version 3.6.1) based on known or predicted associations between these genes. Importantly, we incorporated the corresponding Gene Ontology (GO) terms for each network node, facilitating associated functional annotations. Another network was generated by Cytoscape to establish associations between *mecA* gene expression, antibiotic inhibition diameter and the corresponding phenotype.

#### 1.3.4. Statistical analysis

Data were analyzed using two-way ANOVAs with the genes studied and *S. aureus* strains (*mecA*+ and *mecA*-) as the two predictor variables. Differences at Tukey’s test HSD P < 0.05 were considered statistically significant. Analyses were performed using GraphPad Software (version 6.0, CA, USA). A heatmap was generated to visualize the correlation of the expression of candidate genes in the different strains of *S. aureus* studied based on Pearson’s correlation. Only the comparisons with P < 0.05 were regarded as showing differential expression. The neutral/middle expression was set as the median of all the Ct values from tested varieties; the red color was used to indicate an increase with a Ct value below the median, and the green color indicated a decrease with a Ct value above the median. Analyses were performed with ExpressionSuite™ Software (Applied Biosystems, USA).

## 2. Results

### 2.1. Bacterial isolates

The 30 *S. aureus* strains were recovered from clinical samples obtained essentially from the intensive care unit (Table III). The mean age of the patients was 52 years. Twenty-six patients were male, and four were female (sex ratio =0.86). The distribution analysis showed that most isolates (83.33%) were responsible for invasive infections (bacteremia, deep suppurations, etc.) (Table III).

### 2.2. Antibiotic resistance profile

All isolates were resistant to penicillin G (100%). Three were MRSA (13.33%), 36.66% were resistant to gentamicin, 23.33% to erythromycin, 6.66% to ofloxacin, and 3.33% (01 strain) to teicoplanin. Note that all strains were sensitive to vancomycin **[see Supporting Information Figure S1]**.

### 2.3. Molecular detection

#### 2.3.1. PCR amplification of the *mecA* gene

Detection of the *mecA* gene revealed that 12 strains carried the *mecA* gene. The PCR-amplified DNA products of this gene are shown in **Supporting Information Figure S2A**. These strains are mostly invasive (Table I). Note that only 3 MRSA strains were identified by the phenotypic method and that one did not harbor the *mecA* gene (blood culture) (lane 22 of the agarose gel).

#### 2.3.2. PCR amplification *of the RNAIII* regulatory gene

The results of the amplification of the *RNAIII* regulatory gene allowed us to reveal the presence of a single amplification profile. This is a 250 bp band indicating the presence of *RNAIII* in all strains tested [**Supporting Information Figure S2B**].

#### 2.3.3. PCR amplification of <u>virulence</u> <u>gene</u>s (*spA, SpdC* and *atlA*)

The results obtained allowed us to reveal the presence of all the *spA, SpdC* and *atlA* genes in all the strains studied with a single band profile for each of these genes of 300, 139, and 110 bp, respectively [**Supporting Information Figure S3**].

### 2.4. Functional clustering of virulence genes in *S. aureus*

Our GO enrichment network results revealed that the *WalR, atlA* and *spA* genes form a cluster within the “virulence and two-component system” functional group in our analysis (Figure 1). This observation suggests potential functional links and possible coregulation between these genes in Staphylococcus aureus. Furthermore, these genes are also interconnected by coexpression edges, thus reinforcing the hypothesis of coordination in their gene expression. These findings highlight the importance of these genes in the biology of S. aureus, particularly with regard to virulence and gene regulation. However, one particularly intriguing observation deserves special attention in our analysis of the network. While the *WalR, atlA* and *spA* genes share potential functional links and coexpression edges, *spdC* is distinguished not only by its preferential location in the extracellular region but also by its well-established role as a novel virulence factor in Staphylococcus aureus. This observation highlights a subtlety in the regulation and interactions of these genes. While *WalR, atlA* and *spA* could be involved in intracellular regulatory and virulence processes, *spdC* seems to play a more specific role in the extracellular region, potentially associated with functions such as adhesion or interaction with the environment. extracellular. The proximity of *spdC* to *spA* in this region could suggest coordination between these two genes in key extracellular functions, even if they do not directly share the same functional links with other genes in the “virulence and two-component system” cluster.

**Figure 1:**
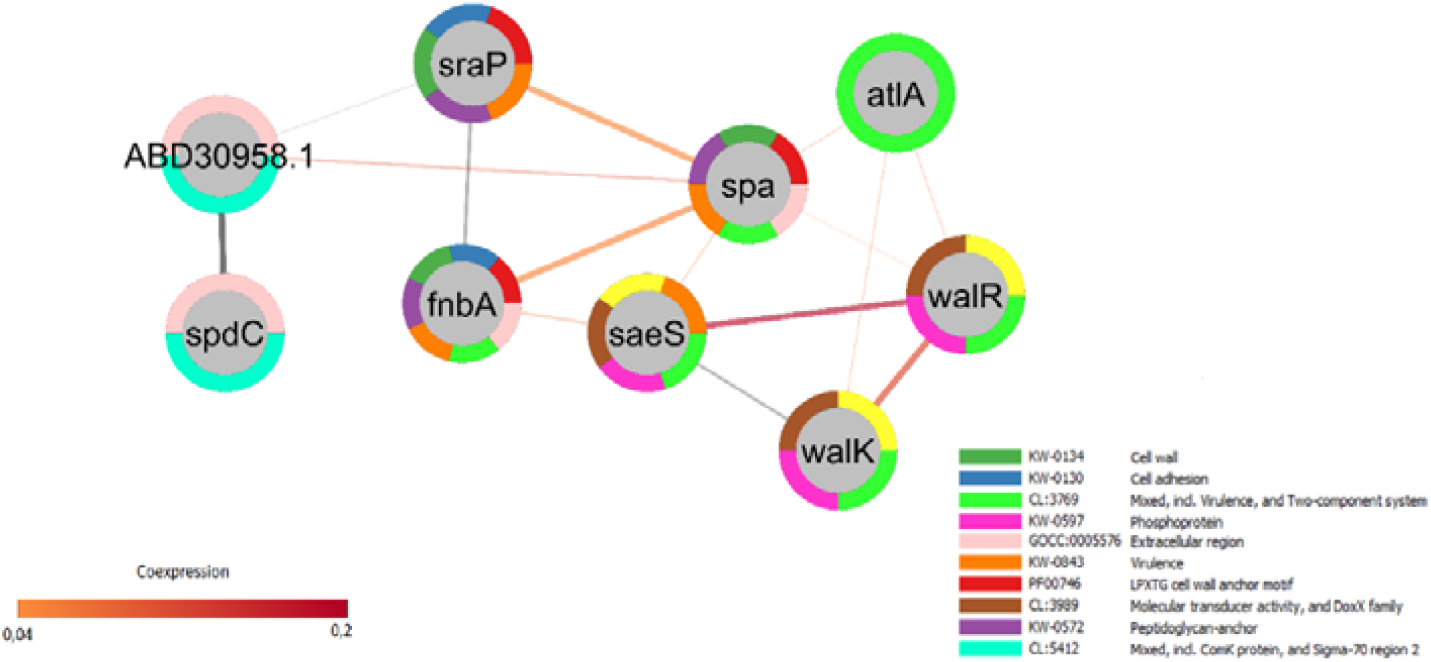
GO Enrichment Network revealing functional links and intriguing localization of virulence genes in *S. aureus*.

### 2.5. Analysis of differentially expressed (*RNAIII, spA, SpdC* and *atlA*) genes by qRT-PCR

The expression profiles of the *RNAIII, spA, SpdC* and *atlA* genes were analyzed from all *S. aureus* strains. Heatmaps of transcript expression were constructed [**see Supporting Information—Figure S4**], and expression comparative analyses were conducted (Figure 2). The results showed a significant accumulation of *RNAIII* transcripts in *mecA*^**-**^ strains compared to *mecA*^**+**^ strains (Figure 1). On the other hand, the *mecA*^**+**^ strains seemed to express the virulence factors more than the *mecA*^**-**^ strains (Figure 2). A two-way ANOVA followed by Tukey’s multiple comparisons tests indicated that relative *RNAIII* expression is statistically significant between *mecA*^**+**^ and *mecA*^**-**^ strains (F1, 11 = 8.929, P= 0.0123). Additionally, the global expression of virulence factors was significantly different between *mecA*^**+**^ and *mecA*^**-**^ strains (F 3, 33 = 25.03, P < 0.0001). With the exception of the other factors, the *SpdC* gene does not vary significantly (P =0.9992). The results also indicated that the interactions between *mecA*^+/-^ strains and virulence factors were significantly different (F3, 33 = 34.02, P < 0.0001).

**Figure 2:**
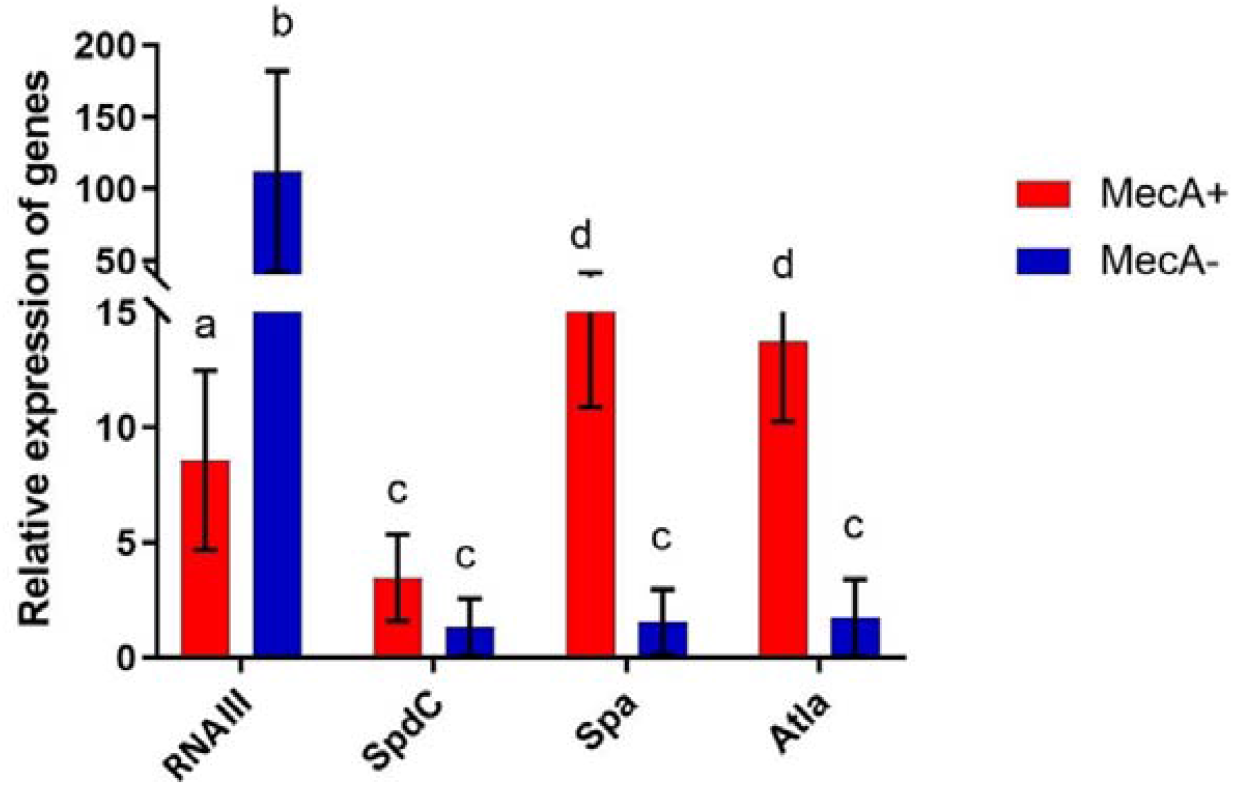
Differential expression of virulence factors (*SpdC, spA* and *atlA*) and regulatory *RNAIII* between *mecA* + and *mecA*-strains. The 16S rRNA gene was used as a reference gene. Error bars show the standard error between three replicates performed. Bars with different letters within each panel are significantly different at *P*□ >□0.05 according to Tukey’s test.

#### 2.5.1. Expression of the *mecA* gene

Twelve *mecA*^**+**^ strains were evaluated for expression of the *mecA* gene (Figure 2). One-way ANOVA followed by Tukey’s multiple comparisons test indicated that relative *mecA* expression was statistically significant between *mecA*^+^ strains (F11, 24= 103.9, P<0.0001).

The results support strong expression of the *mecA* gene in the phenotypically resistant strains S12, S16 and S36. In addition, we observed low expression in other phenotypically sensitive *mecA*^**+**^ strains (Figure 3).

**Figure 3:**
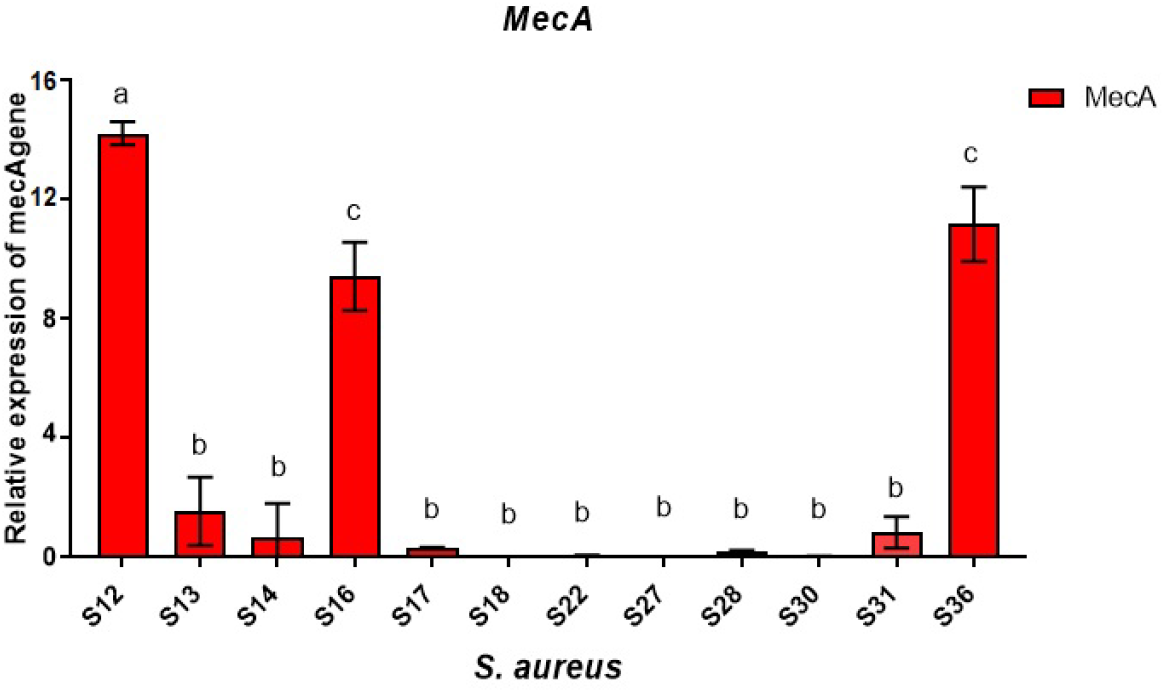
Expression of the *mecA* gene in phenotypically sensitive and methicillin-resistant strains. The 16S rRNA gene was used as a reference gene. Error bars show the standard error between three replicates performed. Bars with different letters within each panel are significantly different at *P*□ >□0.05 according to Tukey’s test.

#### 2.5.2. Correlation between methicillin-resistant/sensitive phenotype and *mecA* expression

To show a correlation between the phenotypic expression of methicillin resistance and the transcription of the *mecA* gene, a multilayer association network was established by cytoscpae 3.0.1 software (Figure 4). In addition, only the strains that are phenotypically resistant (diameter between 1 and 19 mm) to cefoxitin (S12, S16, S36) strongly express the *mecA* gene, whereas phenotypically sensitive strains (diameter between 31 and 38 mm) weakly accumulate *mecA* gene transcripts.

**Figure 4:**
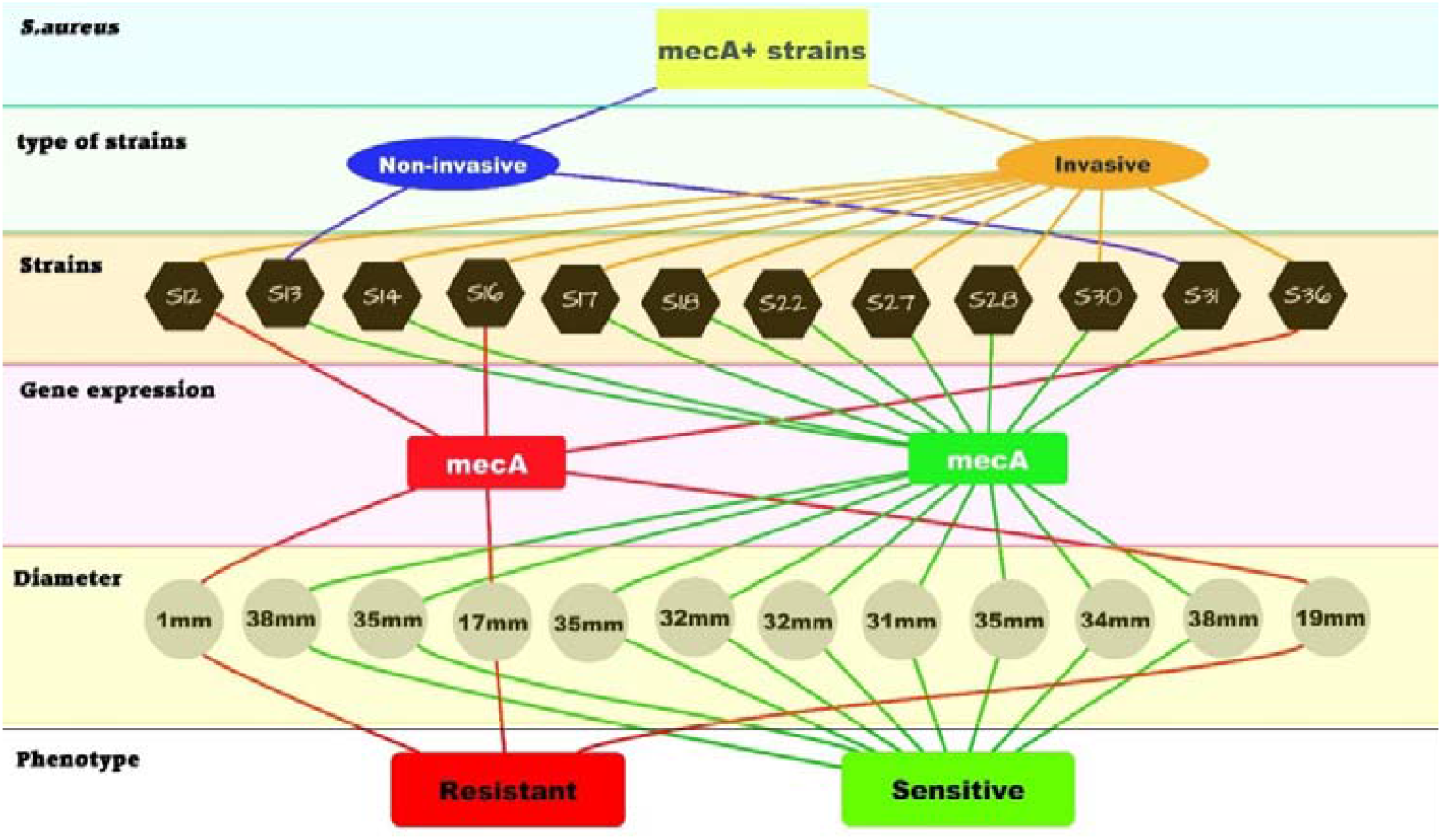
Multilayer association network established by Cytoscape 3.0.1 software

To this end, we determined the Pearson correlation coefficient “r” to demonstrate a statistical correlation between the expression of the *mecA* gene and the diameter of inhibition (Figure 5). This shows a positive correlation “r = 0.8463” between them.

**Figure 5:**
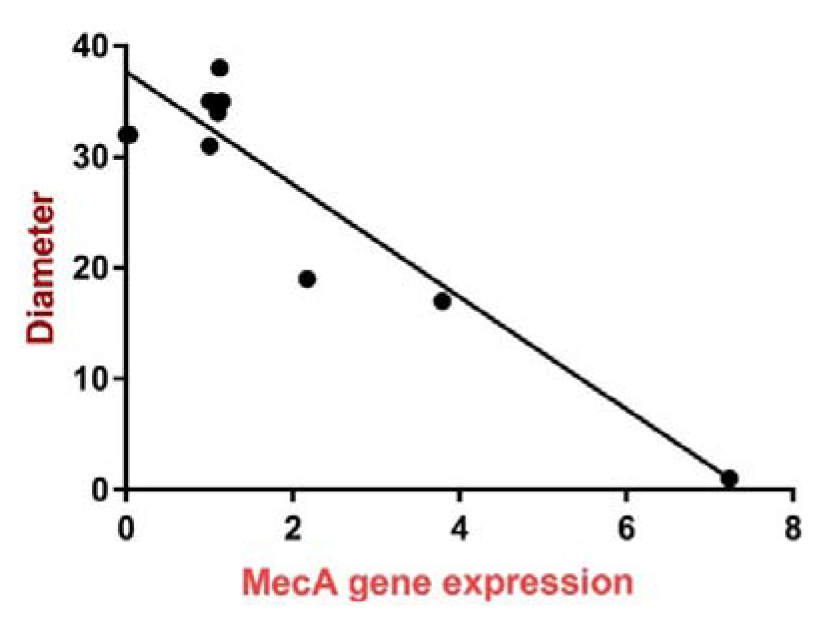
Correlation diagram between *mecA* gene expression and the diameter of the zone of cefoxitin inhibition.

#### 2.5.3. Correlation between the expression of *mecA* and *RNAIII*

By studying the expression of *RNAIII* and the *mecA* gene in *mecA*^**+**^ strains (Figure 6A), our results revealed that the expression of *RNAIII* was higher than the expression of *mecAn* in all strains. This expression was repressed when the strain did not express *mecA* strongly, as in cases S12, S16, and S36 (Figure 6A). A two-way ANOVA followed by Tukey’s multiple comparisons test indicated that relative *RNAIII* and *mecA* expression was statistically significant between each strain, except S28 (F11, 22 = 184.7, P< 0.0001). Additionally, the global expression of genes (*RNAIII* and *mecA*) was significantly different between strains (F 1, 2 = 725.3, P= 0.0014). To demonstrate this negative correlation between *RNAIII* and *mecA*, a test on the Pearson correlation coefficient “r” was established (Figure 6B). The figure shows that the *mecA* gene is inversely expressed with respect to the *RNAIII* gene. There is therefore a negative correlation (0.04) between the expression of *RNAIII* and *mecA*.

**Figure 6:**
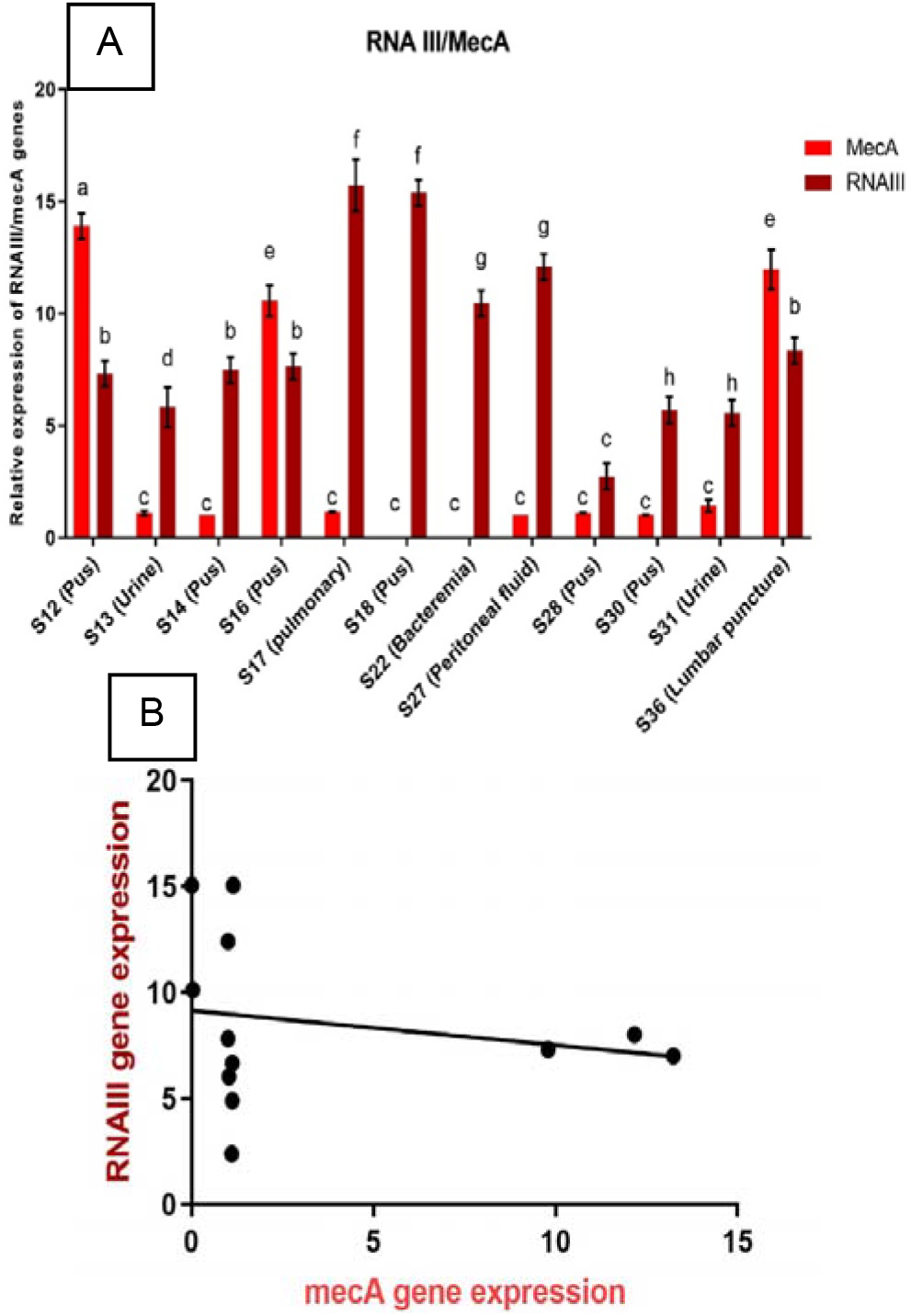
Histograms showing the difference (A) and correlation (B) of gene expression between *RNAIII* and *mecA*

## 3. Discussion

*S. aureus* has an extraordinary repertoire of virulence factors that allows it to survive extreme conditions within the human host [19]. For these bacteria, colonization of the human nose presents a significant challenge that requires not only adherence to nasal epithelial cells but also an ability to cope with host defense and competing resident microorganisms. *S. aureus* adheres to and invades host epithelial cells using a variety of molecules [20]. In response to physiological changes during infection, bacteria can involve factors responsible for the shift of *S. aureus* from commensalism to pathogenesis [21]. One of the factors involved in this passage is a range of virulence factors allowing its adhesion and its invasion [22]. Thirty clinical *S aureus* strains have been collected from different infectious diseases. The presence of all virulence genes (*spdC*, altA, *spA*) and the regulator gene *RNAIII* in invasive and noninvasive strains explains the seriousness of the infections [23]. Among these strains, 70% come from deep suppurations and blood cultures, which thus shows the predominance of isolated invasive strains (83.33%). Indeed, several studies have also found this invasive character, particularly in blood cultures (37%) and in deeply infected skin samples (26.5%) [24].

The rates of methicillin resistance among clinical isolates vary greatly by country, ranging from single-digit rates in Scandinavian countries to over 50%, for example, in the U.S. and China [25]. In Tunisia, this resistance does not exceed 16% (https://www.infectiologie.org.tn/pdf_ppt_docs/resistance/1544644887.pdf). We also showed that 13.33% of the strains isolated were MRSA strains.

Molecular detection of the *mecA* gene in all the isolated strains was carried out. Interestingly, the phenotypic and molecular results between methicillin resistance and the presence of the *mecA* gene revealed that among the 12 *mecA*-positive strains, 9 strains were phenotypically sensitive to methicillin. Indeed, it has been suggested that such isolates could be classified as a new type of MRSA, designated OS-MRSA (*mecA*-positive methicillin-sensitive strains).

These isolates could be misclassified as MSSAn in the daily routine if we base our decisions only on antimicrobial susceptibility testing[26]. As discussed earlier, treating OS-MRSAn infections requires taking precautions because using beta-lactam antibiotics can lead to the emergence of highly resistant MRSA due to the presence of the gene *mecA[26–28]*.

Our results also show that among the four strains of *S. aureus* phenotypically generated with methillin, three were *mecA* positive and one was *mecA* negative. In fact, the absence of the *mecA* gene in strains phenotypically resistant to methicillin suggests other resistance mechanisms apart from the acquisition of the *mecA* gene. Lakhundi & Zhang (2018) discussed homologs of the *mecA* gene (*mecB, mecC, mecD*), which are associated with resistance to beta-lactamins. Another study in Nigeria also reported the complete absence of five main types of SCCmec and *mecA* genes as well as the PLP2a gene product in phenotypically resistant isolates, thus suggesting a likelihood of beta-lactamase hyperproduction as the cause of the phenomenon [30].

All over the world, *S aureus strains* that are phenotypically resistant to methicillin strongly express the *mecA* gene [31]. Our results revealed overexpression of the *mecA* gene in phenotypically resistant strains. In fact, *mecA* transcription and PLP2a expression showed a very positive correlation with methicillin-resistant and methicillin-sensitive phenotypes. The positive relationship between *mecA* transcription levels and methicillin resistance was supported by a previous study [31]. In our study, we investigated the complex interplay between antibiotic resistance and virulence gene expression in *S. aureus*. Our exploration began with the construction of a functional network to better understand the functional associations between virulence genes. Notably, our GO enrichment network analysis revealed intriguing connections between the *WalR, atlA*, and *spA* genes, grouping them within the “virulence and two-component system” functional group. This grouping strongly suggests a potential coregulation between these genes, highlighting their central role in *S. aureus* biology, particularly in terms of virulence and gene regulation, which is consistent with previous research [32,33]. One of the most compelling findings of our study concerns the differential expression of *RNAIII* and virulence factors in *mecA*+ and *mecA*-strains of *S. aureus*. The substantial accumulation of *RNAIII* transcripts in *mecA*-strains compared to *mecA*+ strains reveals a complex regulatory interaction. This suggests that the absence of *mecA*, a key player in methicillin resistance, could influence the expression of virulence factors [34].

Our results reveal that *mecA*-positive strains express more adhesion genes (primarily the *spA* gene) and factors that are involved in biofilm formation (primarily *atlA*), suggesting roles for these proteins in the progression of infection with *S. aureus*, escape to the bloodstream, and colonization of host tissues [35]. These results are supported by Uribe-García et al. [33] who showed that 88% of MRSA strains are biofilm formers. Indeed, the formation of the biofilm allows the persistence of the infection and increases the horizontal transfer of plasmids carrying antibiotic resistance genes [33]. In contrast, *mecA*-negative strains express fewer adhesion factors but overexpress the transcriptional regulator *RNAIII*. This regulator positively regulates proteases and certain toxins [36]. This explains why the strains that do not have the *mecA* gene are more toxic than the positive *mecA* strains. These results were requested by [37], who reported that *mecA* expression affects the *agr* system and reduces toxin production.

On the other hand, qRT-PCR was used to compare the expression of *RNAIII* and *mecA*. The results reveal that there is a negative regulation of gene expression between *RNAIII* and *mecA*, suggesting that the high expression of *mecA* may act on the expression of *RNAIII and suggesting a* possible link between virulence and antibiotic resistance. Indeed, the *mecA* gene has been shown to negatively regulate *RNAIII* [38]. This negative regulation can be explained by the fact that the strong expression of *mecA* (major determinant of resistance to MRSA) can affect the structure of the peptidoglycan or its interaction with other proteins associated with the cell wall and prevent detection of the peptide. Self-inducing (AIP), thus resulting in an insensitive *agr* system and repressing the expression of *RNAIII* [39]. In addition, studies have shown that *agr* dysfunction has been linked to increased biofilm formation and improved colonization capacity [40], which is still consistent with our previous results. The *mecA*-positive strains are more expressive of adhesion factors (*SpA*) and factors contributing to biofilm formation (*atlA*)[38], thus suggesting that these strains are responsible for the dysfunction of the *agr* system. However, it is important to highlight that we did not detect a significant di*spA*rity in the expression level of the *spdC* gene between *mecA*+ and *mecA*-strains. These findings align with those proposed by Bakr et al. [41] who observed a lack of significant correlation between the expression level of the *spdC* gene and the activity of the tested virulence factors or the antimicrobial resistance phenotype.

Along with our results on dual *mecA* activity (antibiotic resistance and virulence regulation), a link between antibiotic resistance and virulence regulation is believed to play an even more important role in pathogenicity. The relationship between virulence and resistance was also noted by Seidl et al. [32], wherein the authors showed that the intrinsic virulence of MRSA strains is similar to, or even less than, that of MSSA and that the increase in virulence is associated with the decrease in methicillin resistance levels [32]. Indeed, the different expression profiles of the genes involved in membership, invasion, and evasion of the immune system, in combination with multidrug resistance to antibiotics, increase the degree of pathogenicity and the spread of infection [33].

## 4. Conclusion

In this manuscript, it was shown that virulence gene expression was higher in MRSA than in MSSA, and we showed a negative correlation between *RNAIII* and *mecA*. An understanding of virulence and resistance relationships would help reduce the impact of *S. aureus* infections. Further studies are needed to provide more detailed information about the different mechanisms of virulence and resistance used by this pathogen and the interaction between them. In addition to many open mechanistic questions, the main current challenge remains how to integrate the gained understanding for the development of anti-virulence drugs or even probiotic approaches for anti-virulence strategies against *S. aureus*.

## Acknowledgements

This work was supported by the Ministry of Higher Education and Scientific Research of Tunisia.

