## Supporting Information Figures for "Impact of methicillin resistance on virulence factor expression in *Staphylococcus aureus*: Insights from gene expression profiling"

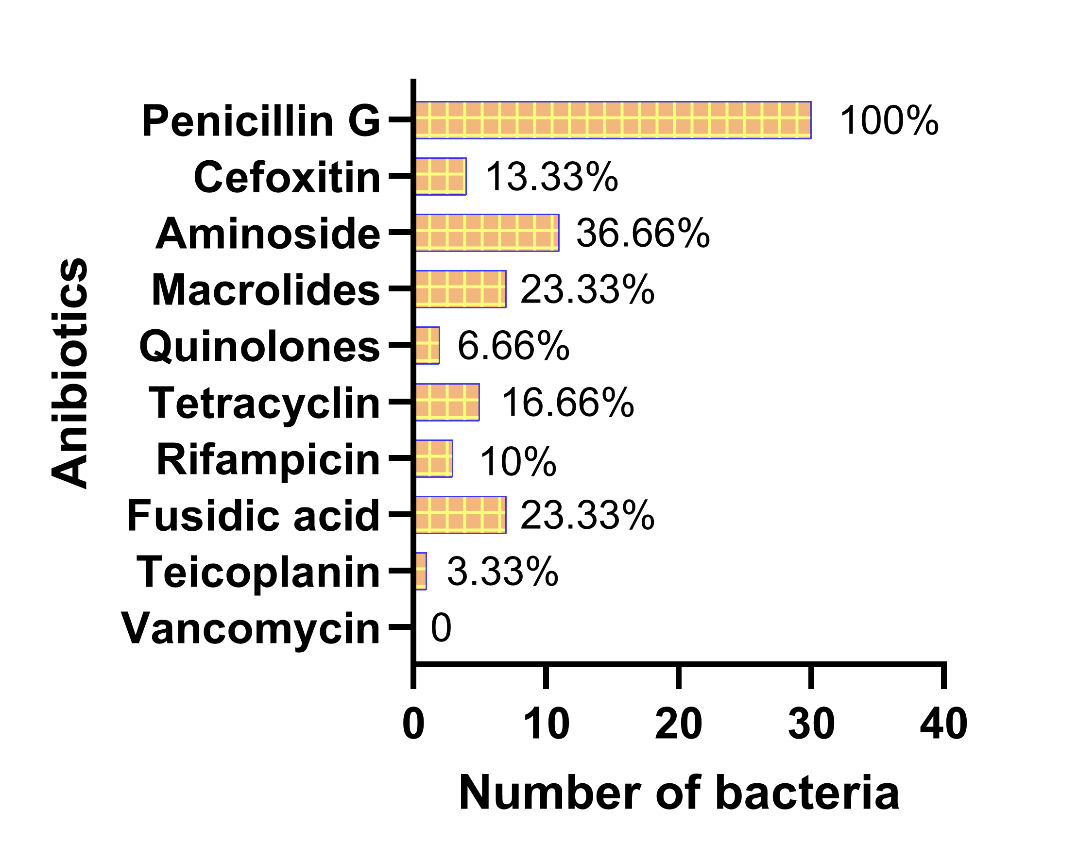

**Figure S1:** Resistance to different anibiotics in *S. aureus* isolates

A

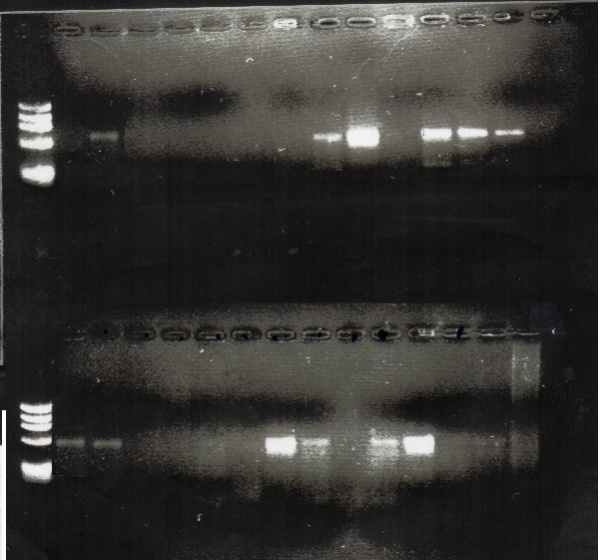

*mecA*

M	1	2	3	4	5	6	7	8	9	10	11 12 13 14 15

M	1	2	3	4	5	6	7	8	9	10	11 12 13 14 15

M 16	17 18 19 20 21 22 23 24 25 26 27 28 29 30

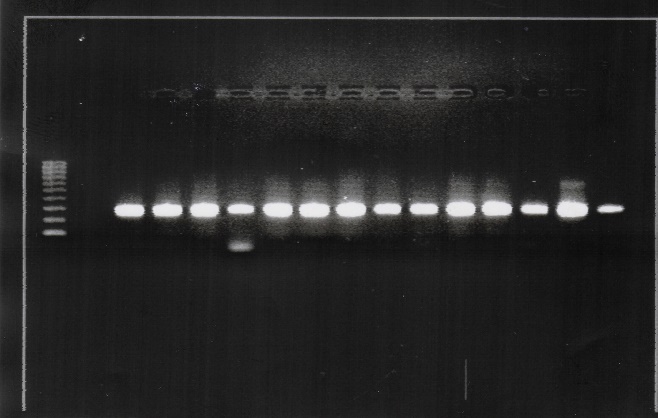

*mecA*

*mecA*

*RNAIII*

**Figure S2:** Representative examples of two agarose gels showing amplifications of ~ 500bp and ~ 250bp and corresponding respectively to regions of the mecA and RNAIII genes. M = 100bp DNA marker fragment; lane (1): Negative control. A: Lanes 2, 8, 9, 11, 12, 13, 16, 17, 22, 23, 25 and 26 indicate mecA + strains while the other lanes indicate mecA- strains. B: all the lanes indicate the presence of the RNAIII gene.

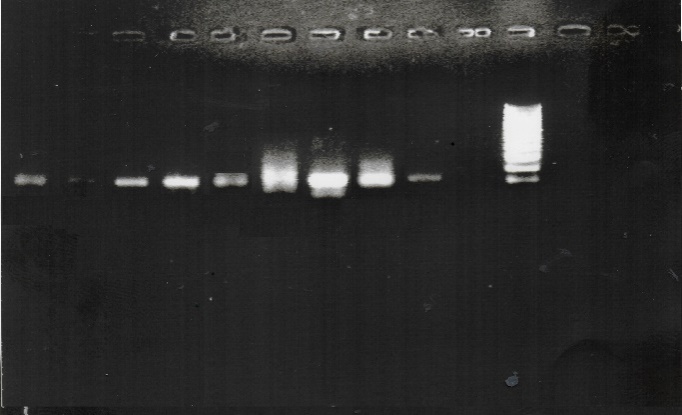

M	1	2	3	4	5	6	7	8	9	10	11	12 13	14 15

M	1	2	3	4	5	6	7	8	9	10

10	9	8	7	6	5	4	3	2	1	M

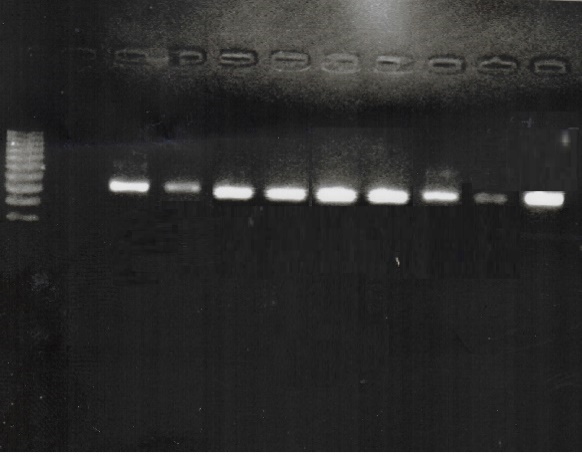

*spA*

*atlA*

*atlA*

*spA*

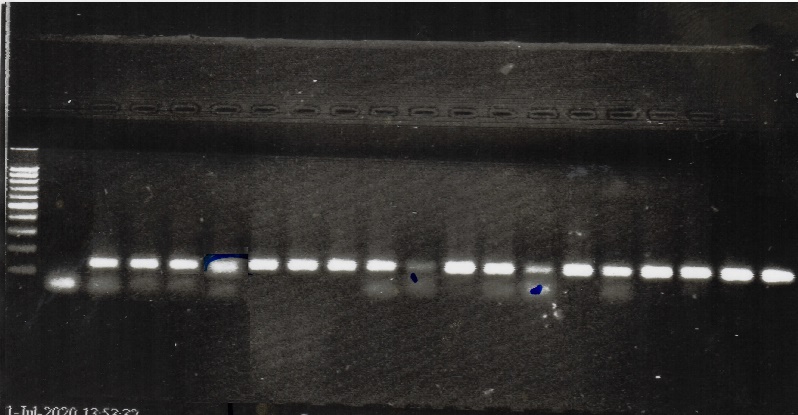

*SpdC*

**Figure S3:** Representative examples of three agarose gel showing amplifications of ~ 110bp, ~ 300bp and ~ 139bp and corresponding respectively to regions of the atlA, spA and SpdC genes. M = 100bp DNA marker fragment; lane (1): Negative control; the other lanes indicate positive strains for spA, SpdC and atlA.

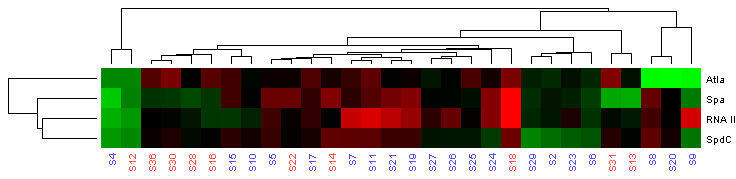

**Figure S4:** Expression profiles of candidate genes. The results of the relative expression levels (Δct) of candidate genes were used for hierarchical cluster analysis with Data Assist v3.01. The colour scale represents relative expression levels, with red as increased transcript abundance and green as decreased transcript abundance.
