## Supporting Information Table for "Impact of methicillin resistance on virulence factor expression in *Staphylococcus aureus*: Insights from gene expression profiling"

**Table S1:** Clinical data of patients infected with *S. aureus*

| Strains | Age | Sex | Type of sample | Antecedents | Antibiotherapy | Phenotype |
| --- | --- | --- | --- | --- | --- | --- |
| S2 | 74 years | women | Cutaneous(superficial pus) | - | - | S |
| S4 | 49 years | man | Cutaneous(superficial pus) | - | - | S |
| S5 | 64 years | man | Blood culture | - | - | S |
| S6 | 66 years | man | Vascular catheter | - | - | S |
| S7 | 65 years | man | Deep suppuration | - | - | S |
| S8 | 57 years | man | Deep suppuration | diabetes and lung cancer | One month of antibiotherapy: ofloxacine (Quinilones ) | S |
| S9 | 74 years | man | Deep suppuration | - | - | S |
| S10 | 21 years | man | Deep suppuration | - | - | S |
| S11 | 37 years | man | LCR | - | - | S |
| S12 | 32 years | women | Peritoneal pus | peritonitis | Bactrim and gentamycin | R |
| S13 | 61 years | man | Urine | - | - | S |
| S14 | 52 years | man | Deep suppuration | - | - | S |
| S15 | 62 years | man | Blood culture |  |  | S |
| S16 | 70 years | man | Peritoneal pus | acute intestinal obstruction | - | R |
| S17 | 33 years | man | pulmonary | - | - | S |
| S18 | 39 years | man | Deep suppuration | - | - | S |
| S19 | 49 years | man | Deep suppuration | - | - | S |
| S20 | 50 years | man | urine | - | - | S |
| S21 | 49 years | man | Blood culture | - | - | S |
| S22 | 56 years | man | Blood culture | Fever and pain fever syndrome left pelvis | *Augmentin (12 days)  *Genta (3 days)  After 12 days: Bactrim +oxacillin) | S |
| S23 | 66 years | women | Blood culture | - | - | S |
| S24 | 65 years | man | Blood culture | - | - | S |
| S25 | 62 ans | Man | Blood culture | -Central venous catheter removal  - bacteremia linked to catheter | -Claforan (céfoxtaxim)  - 3 days vanco-teico | S |
| S26 | 67 years | man | Blood culture | - | - | S |
| S27 | 20 days | man | Periotoneal liquid | Kidney cyst | Under peritonitis : Vanco-fortum | S |
| S28 | 62 years | man | catheter | Renal insufficiency | rifocin | S |
| S29 | 28 jours | premature | Blood culture | Respiratory distress | - | R |
| S30 | 44 years | man | Deep suppuration | - | - | S |
| S31 | 61 years | man | Urine | - | - | S |
| S36 | 45 years | man | Pleural liquid | - | - | R |

**Table S2 :** List of primersused for PCR and qRT-PCR analysis

| **Genes** | **Primer sequences**  **5’🡺3’** | **Amplicon size** | **References** |
| --- | --- | --- | --- |
| ***spA*** | F:AAGTGCTAACCTATTGTCAGAAG  R:TCGTCTTTAAGGCTTTGGATG | **300pb** | (Poupel et al. 2018) |
| ***SpdC*** | F:GCAGTAGGATACATTGGTT  R:CAGCCTCAGTATGATTAGTT | **139pb** |  |
| ***atlA*** | F:AACAGCCACCAACGGATTAC  R:CATAGTCAGCATAGTTATTCATTG | **110pb** |  |
| ***RNAIII*** | F : CCTAGATCACAGAGATGTGATGGR: AATACATAGCACTGAGTCCAAGG | **250pb** | (Yan et al. 2015) |
| ***mecA*** | F : AAAATCGATGGTAAAGGTTGGCR R:AGTTCTGCAGTACCGCATTTG | **530pb** | (Shanehbandi et al. 2014) |
